# Disordered proteins: microphases or associative polymers?

**DOI:** 10.1101/2024.10.09.617362

**Authors:** Martin Girard

## Abstract

We develop a surrogate model for low complexity disordered proteins, which allows us to generate sequences with quantifiable disorder. We investigate properties of these sequences, and show that the sequence dependence of the radius of gyration only arises in the vicinity of the polymer collapse transition. Microphase propensity of the sequence is shown to be a reliable predictor, outperforming state of the art methods, in the crossover region. We show that predictions of associative polymer theory arises only as a limiting case, and discuss its applicability.

## MAIN

Liquid-liquid phase separation has emerged as a topic of great importance in molecular biology [1], with particular attention given to intrinsically disordered proteins (IDPs). A great deal of effort has been dedicated to understanding the relation between protein sequence and phase separation, often referred to as molecular grammar [2–4]. On the theoretical side, associative polymers have emerged as a model to understand properties and phase behavior of disordered proteins [5–11]. Applicability of this model has been recently discussed in [12], where authors found that weak collective interactions are generally more compatible with the liquid phases observed in experiments, as opposed to few strong stickers. However, the field has been limited by the availability of suitable polymer models to characterize sequence disorder.

Mathematical tools from combinatorics provide an ideal framework to label, and describe complexity of sequences. The particular branch dedicated to this is called combinatorics on words. We refer to well established literature for a thorough formal description of the field [13–15]. In this article, we use notions inherited from combinatorics on words, namely balance, to quantify disorder [13]. By using cutting sequence constructions, we create sequences with desired disorder. By analogy to the low-complexity Sturmian sequences, we term these polymers Sturmian polymers [16]. We show that for heteropolymers, sequence dependence only arises in the vicinity of the collapse transition. Within this region, polymer properties such as the radius of gyration can be predicted by characterizing microphase propensity of the sequence, which outperforms current state of the art metrics such as sequence hydropathy decoration [17].

For Sturmian polymers, theoretical prediction made for associative polymers in [18] only arise as a particular limit of our generic model, which is unlikely to arise in the context of low complexity proteins. We also investigate percolation and clustering in these systems. Specifically, we find finite size clusters, as in [19]. We argue that they arise due to finite size effects near the critical temperature, as in [20], and should be instead understood in a finite size scaling context.

### Sturmian polymers as a surrogate model

In order to establish a common language in the context of proteins and polymers, we introduce basic concepts here. We define a word *w* as any sequence of characters, which may be infinite. The set of characters allowed in the sequence constitute the alphabet 𝒜. In the context of proteins, the alphabet consist of letters associated to amino acids : 𝒜 : {*A, C, D*, …, *Y* }. It is often convenient to use an algebraic formulation in terms of *factors*, where the product operation represents concatenation, e.g. *w* = *u* · *v*, where *u* and *v* are also words. The term subword is also used in the literature, which can be used interchangeably with *factors*. To maintain consistency with polymer literature, we use *N* to denote the length of full word, and *n* the length of factors, with obviously *n* ≤ *N*. Words can be characterized by a complexity function *P*_*n*_(*w*), which is equal to the number of unique factors of *w* of length *n*.

The relevant quantity in the context of polymers is balance. Namely a word *w* is said to be ℵ-balanced iff for any two factors of equal length *u, v* of *w*, the number of occurrences of any letter in each of the two factors differs by at most ℵ. We also note that the balance of a factor *u* of *w* is at most the balance of *w*. From the point of view of polymer physics, this quantity is a measure of the largest “block” along the chain. The notion of “block” here being mathematical, we show a few concrete examples of sequences and their balance on Tab I. The product of two factors, identical or not, of the same balance always yield a balance equal or greater than any of the two factors. We label a factor as idempotent if bal(*w*) = bal(*w*^2^).

**Table I.** Examples of binary sequences and their balance.

| Sequence | Balance |
| --- | --- |
| $(AB)^n$ | 1 |
| $(AABB)^n$ | 2 |
| $(A^p B^q)^n$ | $\min(p, q)$ |
| $AABB(A^p B^q)^n$ | $\max(2, \min(p, q))$ |

While the relevant alphabet for biology is of size |𝒜| = 20, for the sake of simplicity, we limit ourselves for the moment to binary alphabets 𝒜 : {*A, B*}, and tackle the properties of *infinite* words. For such words, the complexity function can reveal a lot of details about the properties of the sequence; for instance, any word *w* with P_*n*_(w) < *n* + 1 for any *n* ≥ 2 is ultimately periodic. Conversely, the minimal complexity of an aperiodic binary word is P_*n*_(*w*) = *n* + 1. Aperiodic words of minimal complexity are termed Sturmian words, and their properties are well characterized [16]. The relevant quantity for this article is that all Sturmian words are 1-balanced. We refer the reader to the established literature for more details.

We now turn our attention to *d*-dimensional hypercubic billiard words [21, 22]. These can be constructed using a cutting sequence, i.e. the intersection of lines with square grids [23, 24]. Briefly speaking, a line in *d*-dimensional space with slopes 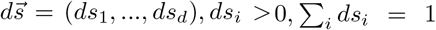 is drawn starting from a fixed point, here taken to be the origin. We now move along this line, so that our position in *d*-dimensional space is 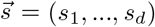. Each time the line intersects a hyperplane corresponding to s_*i*_ = *x*, such that *x* ∈ ℕ, we add the character *i* to the sequence. This construction yields an infinite word, with well defined properties; the alphabet size is *d*, the fraction of character *i* is ds_*i*_, the balance of this word is *d* − 1, and the complexity function grows asymptotically P_*n*_(*w*) ∼ n^*d*−1^. Unsurprisingly, the construction for *d* = 2 is a Sturmian word.

Finite-sized words are slightly different. We interpret them here as finite factor (subword) of an infinite word. The balance of a finite word cannot be larger than the total number of occurrences in the word. Additionally, the notion of periodicity is inapplicable to a finite sequence. For the sake of simplicity, we will abuse formal mathematical notation in this paper. Namely, we will not distinguish finite words, e.g. Christoffel words, from their infinite counterparts [16]. Furthermore, we will refer to all finite sequences derived from hypercubic billiard sequences of balance ℵ as ℵ-balanced Sturmian words, whether they are full length sequences, projections or obtained by concatenation of multiple words. These should be viewed as words that show similar regularity as hypercubic billiard sequences.

We call Sturmian polymer, any polymer whose sequence is derived from a ℵ-balanced Sturmian word. Within this article, we investigate only binary sequences, comprised of monomers *A* and *B*. This has been extensively used in the theory of sticker and spacers, where one of these monomers represents the “sticky” amino acids. These sticky residues are typically interpreted to be aromatics, i.e. phenylalanine, tyrosine and tryptophan, although the actual definition may be protein dependent [25, 26]. For the sake of this article, stickers can be roughly understood as being mainly driven by hydrophobicity, which is reflected in our force-field representation, *vide infra*. In addition to the balance ℵ, the other relevant sequence parameter is the fraction of stickers *f*. The energy scale is parametrized by an effective mean-field interaction parameter χ_*e*_, which takes into account *f* (see methods). This energy scale can be directly interpreted as typical monomer interactions would have, if the monomers were perfectly mixed. This can be used to derive an effective microphase separation propensity, 𝒲 = ℵ(χ_*AA*_ − χ_*e*_)ε, where χ_*AA*_ is the interactions of stickers and ε the energy unit (see methods). This is empirically formulated as the equivalent to the usual co-block microphase propensity.

### Sturmian polymer model

In the context of this paper, we are interested in the properties of relatively low balanced words, e.g. ℵ ≤ 5. At we aim to understand fundamental differences with homopolymers, we are interested in the properties of relatively simple heteropolymers. We consider here binary ℵ-balanced Sturmian sequences. To construct these, we create (2ℵ − 1)-Sturmian sequences with random slopes, then project the first (last) ℵ dimensions to character A (B). The projection breaks the expected balance of the sequence, and this process is not guaranteed to yield a ℵ-balanced sequence. We therefore repeat the process until the sequence has balance ℵ. In addition to these full length sequences, we also construct short *N* = 32 idempotent sequences, which we concatenate to obtain polymers of arbitrary length and identical balance. Concatomers have the additional property of showing constant *f* independently of *N*, while their behavior is indisguishable from their full length counterparts. Homopolymers (ℵ = 0) are only comprised of A beads. We describe our chains as a simple bead-spring polymers, where monomers are interacting through Ashbaugh-Hatch potentials:

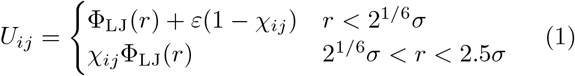

Where ε = 1*k*_*B*_*T* sets the energy scale and Φ_LJ_(r) is the usual Lennard-Jones potential:

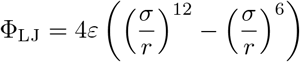

Chain connectivity is enforced by harmonic bonds, where the resting distance (*r*_0_ = 1.1*σ*) is chosen to allow bond crossing, resulting in absence of entanglements. The spring constant is chosen to be *k* = 1000*ϵ*/*σ*^2^. For non-bonded interactions, we use the combination rule 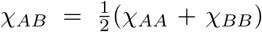. This choice is motivated by two factors. First these potentials are very similar to coarse-grained force-fields for IDPs [27–29]. Second, the effective mean-field parameter is described by a simple average:

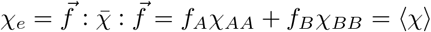

The theta parameter χ_*θ*_ at which an infinite chain undergoes collapse is not yet determined for this potential. For bead-spring chains interacting through Lennard-Jones potentials, the value has been measured to be χ_*θ*_ ≈ 0.22 [30, 31]. At these interaction levels, the Ashbaugh-Hatch potential has increased excluded volume interactions, which yields a higher value of χ_*θ*_. Consistently, we observe good solvent behavior for χ_*e*_ = 0.32, and eventual collapse for χ_*e*_ = 0.45. Unless otherwise noted, we use χ_*BB*_ = 0.2, which corresponds to spacers in a good solvent.

### Sequence specificity, collapse transition and microphase propensity are related

We first investigate behavior of single chains, i.e. in the infinite dilution limit, and focus on the radius of gyration, *R*_*g*_. It is useful to first review classical theory for homopolymers [32]. In the asymptotic limit *N* → ∞, theory predicts a scaling *R*_*g*_ ∼ *N* ^*v*^, where *v* is the so-called polymer exponent. For polymers in a poor solvent, attraction dominates, chains eventually collapse and one obtains *v* = 1/3. Conversely, for polymers in a good solvent, excluded volume wins, and *v* = 0.588. At the theta temperature, attraction exactly compensate excluded volume, chains are ideal, and *v* = 1/2. For finite chains, *R*_*g*_ will exhibit three distinct regimes depending on *N* [32]. There are two length scales to consider, the persistence length 𝓁, and the thermal blob size *g*. The first is induced by the backbone stiffness induced by local interactions along the discrete bonds. Consequently we have 𝓁 ∼ 𝒪(1), and *R*_*g*_ ∼ *N* for *N* ≪ 𝓁. The second is linked to the competition between entropy and enthalpy. In short chains, attractions are weak and unable to collapse the chain, which displays ideal conformations. The scaling *R*_*g*_ ∼ *N* ^1*/*2^ is therefore observed for 𝓁 ≪ *N* ≪ *g*. The transition from *R*_*g*_ ∼ *N* ^1*/*2^ to *R*_*g*_ ∼ *N* ^1*/*3^ is a crossover regime, which can be used to determine *g*. The reader should also note that *N* > *g* is a *sine qua non* condition for multi chain systems to undergo phase separation into polymer-rich and polymer-poor phases.

We now examine simulations of single chains. The radius of gyration exhibited by Sturmian chains is close to homopolymers with equal χ_*e*_ (see Fig 1A). Very little deviation from the homopolymer chains is observed for both short and long chains. However, in the poor solvent crossover regime *N* ≈ g, large deviations from homopolymer are observed. Sequence specificity for disordered chains only appear in this crossover regime. In other words, operating in the regime *N* ∼ *g* amplifies the sequence specificity, and it is likely of functional relevance to cells. It is tempting to assume that this dependence is linked to a sequence dependence of the thermal blob size; however, the picture of thermal blobs as used in a scaling approach is only valid if *N* ≫ *g*.

**Figure 1.**
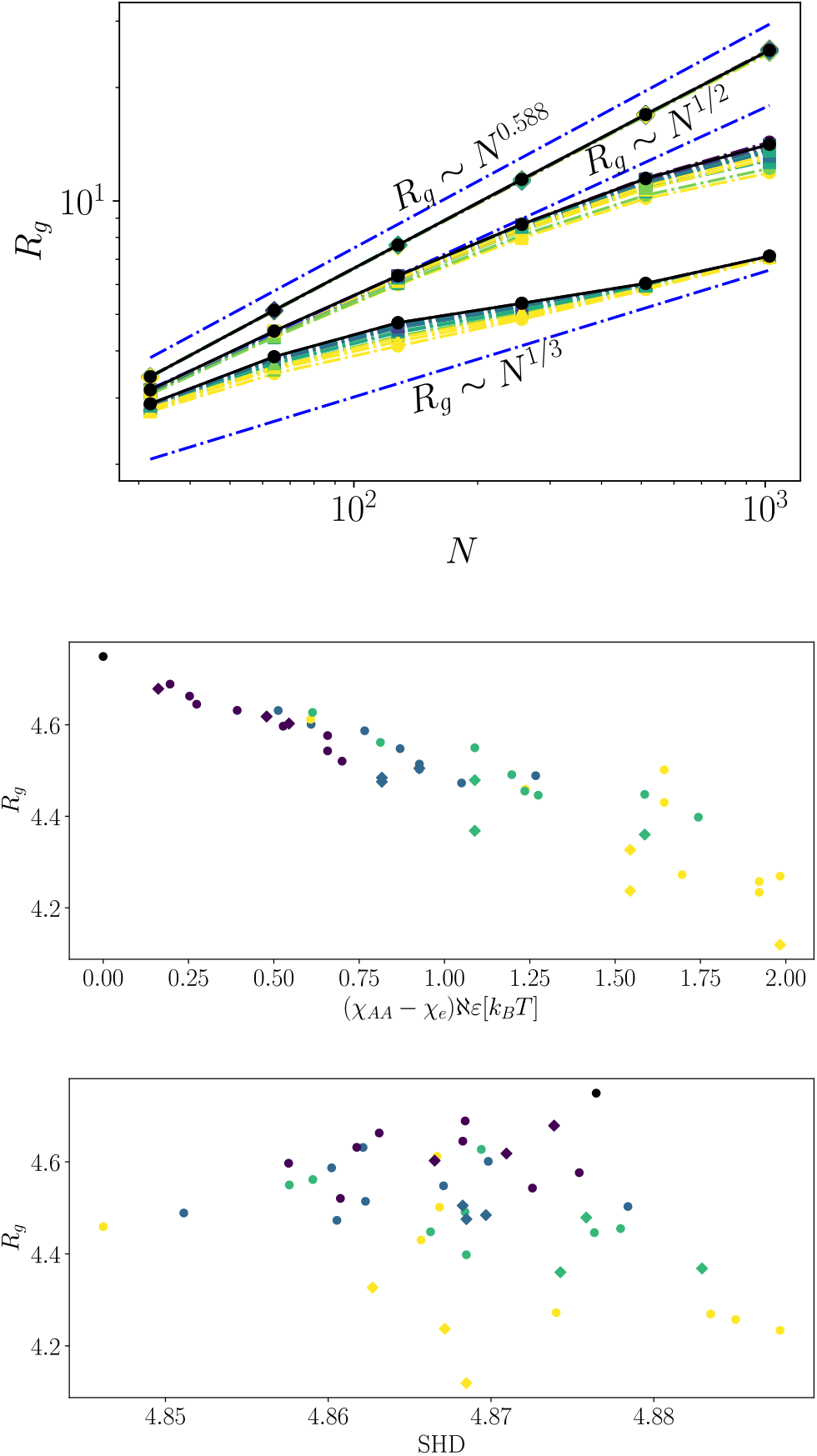
Radius of gyration of Sturmian binary polymers (homopolymer in black). Error bars are smaller than symbol size. Distribution of fraction of stickers *f* depends on ℵ (see SI). Color indicates balance; homopolymers in black, and Sturmian balance ranging from 1 (blue) to 5 (yellow) A) Radius of gyration versus polymer length for *χ*_*e*_ = 0.32, 0.45, 0.55. Scaling relations for *N*^*v*^ in good solvent (*v* = 0.588), theta-solvent (*v* = 1*/*2) and poor solvent (*v* = 1*/*3) are shown as guide for the eye. Upper line is *χ*_*e*_ = 0.32, which follows good solvent scaling. Short chains show ideal scaling, as expected from usual polymer theory. Deviations from homopolymer scaling is only observed near the departure from ideal polymers. B) Variation of *R*_*g*_ versus microphase propensity for *N* = 128, *χ*_*e*_ = 0.55 showing increasing deviation from homopolymers with increasing ℵ. C) Correlation of *R*_*g*_ with state-of-the art sequence hydropathy decoration parameter doesn’t show predictive power over such minor differences. Symbols in B, C indicate whether the chain is a concatomer (diamond) or full length Sturmian polymer (circle).

The deviation of Sturmian polymers from homopolymer directly correlates with the microphase propensity 𝒲 (see Fig 1B), with a nearly linear relation. This takes place in a regime where state of the art predictors, such as sequence hydropathy decoration (SHD) [17], are poor predictors (see Fig 1C). This is likely due to the low complexity of our sequences. Here, the SHD span values between ≈ 4.85 to ≈ 4.88. The sequences considered in [17] show much higher complexity, with SHD spanning values between ≈ 4 and ≈ 7. The parameter 𝒲 should also not be interpreted as a general predictor of *R*_*g*_, as the correlation highly depends on chains being of lengths commensurate with *g*. This hints at partial collapse of some polymer segments before the rest of the chain, altering phase behavior in the vicinity of the critical point.

### Associative polymer theory is an unrealistic limiting case

We now turn to phase separation, and melt properties of Sturmian polymers. We characterize melts by their mean density ⟨*ρ*⟩, computed at the critical Flory-Huggins pressure of a homopolymer *P* ^∗^*σ*^3^/*ε* = *N*/2 − *N* ^−1*/*2^ − log(1 −(1 +*N* ^1*/*2^)^−1^) [33]. Phase separation is characterized by a critical interaction χ_*c*_ = 1/*T*_*c*_, where *T*_*c*_ is the critical temperature, measured by slab simulations (see Methods and SI). The underlying theory for associative polymer [18] is derived in the limit of small *f*, and makes several important predictions. First, phase separation is only determined by sticker strength χ_*AA*_. Second, melt density is independent of χ_*e*_.

We first consider the case of athermal spacers, χ_*BB*_ = 0. Density isolines in the (*f*, χ_*e*_) plane take a hyperbolic appearance (see Fig 2A); For large f, they are parallel to the *f* axis, whereas for large χ_*e*_, they become parallel to the χ_*e*_ axis. There is therefore a regime, in which associative polymer behavior (χ_*e*_-independent density) is reached. This is the associative polymer limit, which is only reached here for very strong interactions, and in the limit of *f* → 0. We obtain similar results for phase separation (see SI), where our lowest value considered, *f* = 0.08, is still too large to be in the associative polymer regime.

**Figure 2.**
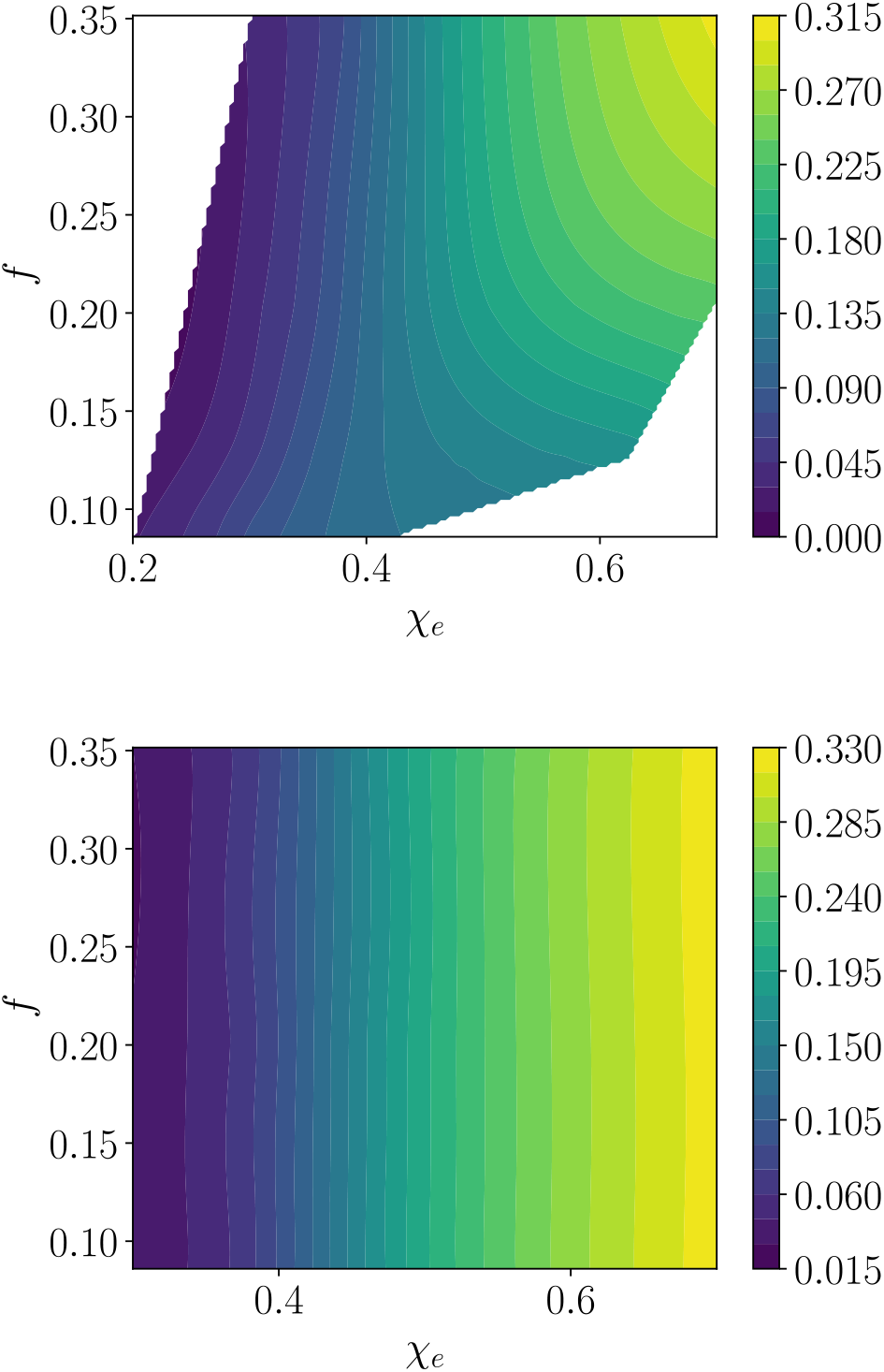
Mean densities obtained for ℵ = 1 binary Sturmian polymer melts for (top)*χ*_*BB*_ = 0 and (bottom) *χ*_*BB*_ = *χ*_*AA*_*/*3. Range of values considered for the top figure is limited by equilibration issues (see Methods)

Spacers, typically the non-aromatic residues, are not athermal in nature [34], and consistently typical force-field parametrizations shows significant interactions [27, 28]. Let us briefly consider spacer interactions weaker than typically observed force-fields, with χ_*BB*_ = χ_*AA*_/3. The density isolines for such cases are completely parallel to the *f* axis (see Fig. 2B), indicating that the associative polymer limit is never reached for reasonable parameters. An alternative parametrization for spacers, χ_*BB*_ = 0.2 also shows a suppressed transition to the as sociative polymer regime.

### Clustering and percolation

We now consider clusters in the polymer melts. Since we have constant pressure simulations with periodic boundary conditions, we can take the usual percolation definition; any two beads within a distance *r*_*c*_ = 1.25σ are part of the same cluster. A cluster is defined as percolating, if it forms a single cluster with any of its periodic images. In a trajectory, a system is percolated if more than half of the configurations show a percolating cluster. For all of our polymer melt simulations, the systems are already percolated. This should not be a surprise to the reader since all of these systems are above the polymer overlap concentration c^∗^ [32].

For direct coexistence simulations, the distribution of clusters forms non-exponential curves when χ_*e*_ ≈ χ_*c*_ (see Fig. 3). These are present independently of the polymer architecture. These are similar to the clusters reported in [19], which clearly shows that associative polymers are not required to find finite-size clusters in direct coexistence. Instead, they are likely due to the proximity of χ_*c*_. In order to do so, we need to point out two flaws in the argument presented in the SI of [19]. Invoking classical nucleation theory for a cluster of *n* particles in equilibrium with a dilute solution, and disregarding prefactors, we write the change in free energy due to cluster formation as:

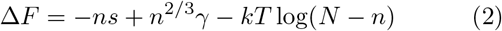

Where s is the supersaturation, and *γ* the interfacial tension. As the cluster grows and the solution becomes dilute, the translational entropy increases. Using an ideal gas approximation, this entropy can be estimated as proportional to log(*c*_*sat*_) ∼ log(*N* − *n*), which was missing from [19]. In addition, the parameters s and *γ* in eq. 2 are generally not constants in the vicinity of χ_*e*_, and are function of (χ_*e*_ − χ_*c*_). For instance, the interfacial tension varies as *γ* ∼ (χ_*e*_ − χ_*c*_)^*µ*^, where µ ≈ 1.26 is a critical exponent [31]. These two additions give rise to a finite fraction of the system in the condensed phase near χ_*e*_ ≈ χ_*c*_, as would be expected from the binodal curve. Cluster size fluctuation increase in the vicinity of χ_*c*_, as expected from critical behavior. This gives produces an illusion of finite-size clusters, which would vanish in an infinite system. In absence of finite size scaling analysis, it is therefore impossible to determine whether the clusters reported in simulations of [19] are relevant for the experimental conditions they are describing.

**Figure 3.**
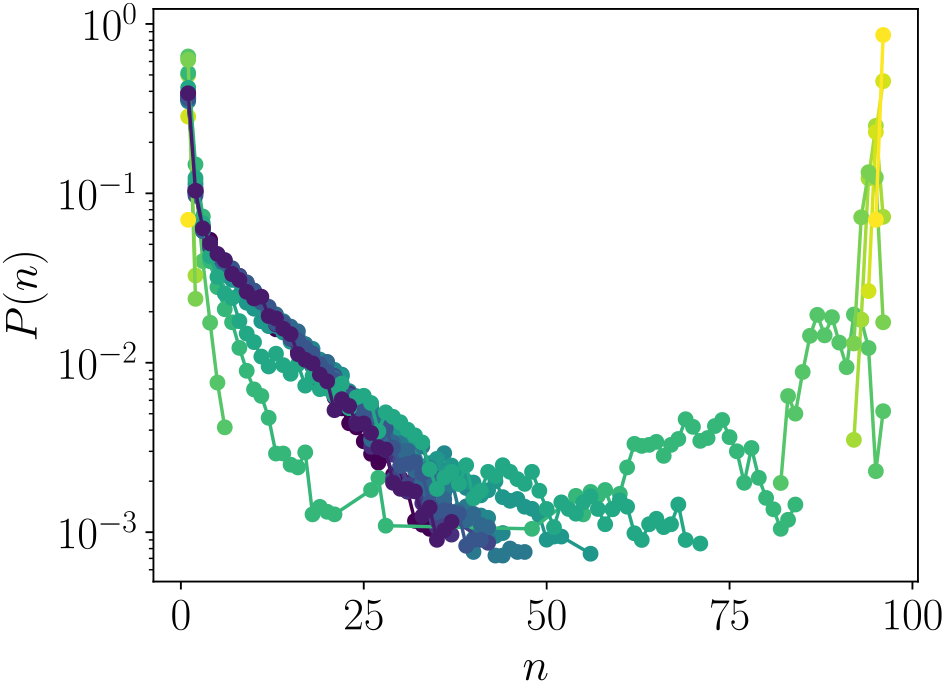
Probability of observing a cluster of size *n, P* (*n*) for *f* ≈ 0.08,ℵ = 1 binary Sturmian polymers with *χ*_*BB*_ = *χ*_*AA*_*/*3. Color indicates the value of *χ*_*e*_. The system undergoes phase separation, which is manifested through finite-size clusters.

## DISCUSSION

Behavior of our polymer system varies depending on the solvent quality of spacers. Associative polymer theory assumes that spacers are athermal. However, usual force-field parametrization yields spacers in good solvent conditions, which suppress the transition towards associative polymers [27, 35]. This thermal parametrization is also supported experimentally in [34]. Taken together, these results support the results of [12].

The polymer model described here is a systematic binary approach for heteropolymers, with hydropathy patterning. The cutting word procedure can be straightforwardly extended to include charges. This provides a clear pathway to investigate the role of disorder in polyam-pholytes, and its relation to coacervation [36]; which would closely mirror our approach for hydrophobicity.

Microphase propensity being an excellent predictor of polymer properties is rather surprising. We conjecture that this is linked to the collapse of segments enriched in stickers, leading to the whole chain collapsing. Such a parameter would normally be associated with the formation of microstructures in the dense phase. This would be consistent with the picture previously associated with percolation in the dense phase [7]. In the co-block polymer literature, the threshold for structure formation is usually estimated to be 𝒲 ∼ 10 kT [37, 38], but can be significantly larger for asymmetric (*f*≠ 1/2) polymers. It is therefore unlikely that low complexity sequences are able to undergo microphase separation on their own. However, this does not preclude that post-translational modifications can alter the microphase propensity.

Our surrogate model predicts that sequence only plays a role near the polymer collapse transition. For homopolymers, this represents a thermodynamic transition in the limit *N* → ∞, in which susceptibilities diverge. It would be tempting to associate a functional role to sequence properties here, such as protein recognition. However, the reader should note that proteins are often modified post-translationally, for instance through methylation. For efficiency reasons, it is likely that evolution has designed proteins to show large response to a few modifications. As with sequence disorder, this would also arise through the vicinity of a collapse transition. These functional aspects therefore need to be further investigated.

## METHODS

To determine single chain properties, we run simulations of 64 independent chains for each realization of the sequence, using the HOOMD-Blue molecular dynamics package [39–42].

We use a timestep Δ*t* = 0.01*τ*, where *τ* = *σ*ℳ^1*/*2^/*ε*^1*/*2^ is the natural time unit of the system. Simulations run for t = 5 · 10^5^τ. All beads have an equal mass of M. A Langevin thermostat with friction constant *γ* = 0.10M/τ is applied to obtain a canonical ensemble.

For each value of ℵ > 0, we generate 8 distinct sequences. The slopes of the cutting word are chosen randomly (uniform distribution). The sticker fraction is a by-product of the slopes, and is therefore also randomly distributed. The resulting distribution of *f* peaks near *f* = 1/2, and becomes sharper for larger ℵ. We generate 3 concatomers for each value of ℵ. Taken together, this results in simulation of 1026 independent simulations, each of 64 chains. For simulations of melts and phase behavior, we use simulations with 96 chains. Large values of χ_*e*_ coupled to small values of *f* leads to large sticker attractions. We choose state points such that χ_*AA*_ ≤ 5. Simulations with different χ_*e*_ are coupled by Hamiltonian replica exchange to avoid slow relaxation times, with exchanges attempted every 3 · 10^4^ timesteps. This is required to avoid slow relaxation times when χ_*AA*_ ≈ 5. For melts (direct coexistance), for each value of *f*, we use 40 (16) values of χ_*e*_, linearly spaced. The range of χ_*e*_ value considered yields an acceptance rate ≳ 15% for all replicas. Simulations are run for *t* = 1.25 · 10^6^*τ*. For melts, we use a Bussi thermostat [43] with relaxation *τ* = 5, and a Langevin piston [44] with relaxation constant *τ*_*p*_ = 20, friction coefficient *γ*_*P*_ = 1 and cubic pressure coupling. For direct coexistance simulations, we use an hexagonal prism geometry, with side length 25σ, and mean density ⟨ρ⟩ = 0.05. Unless otherwise specified, integration parameters are otherwise identical to single chains.

Cutting words are created using a custom python package gitlab.mpcdf.mpg.de/mgirard/Words, which also provides the sequence hydropathy decoration analysis. The version used for the current article is also available as supplementary data. Molecular topologies are assembled using hoobas [45]. Analysis is performed using the freud package [46]. Computational workflow was managed by the signac-flow package [47–49].

## ACKNOWLEDGEMENTS

I am grateful for numerous important discussions on polymer theory with Kurt Kremer and Burkhard Dünweg, and on disordered proteins with Edward A. Lemke. I thank Arash Nikoubashman, Lukas Stelzl, Edward A. Lemke, and Kurt Kremer for a critical reading of this manuscript.

This work was supported by the Max-Planck Computing and Data Facilities, and financial support of the Collaborative Research Centre 1551 “Polymer concepts in cellular function” of the DFG (Deutsche Forschungsge-meinschaft, project number 464588647).

