## Supplemental information for "Disordered proteins: microphases or associative polymers?"

### Supplementary information

Martin Girard*
*Max Planck Institute for Polymer Research, Ackermannweg 10, Mainz 55128, Germany and Institute for Quantitative and Computational Biosciences, Johannes-von-Müller-Weg 6, Mainz 55128, Germany*

#### MELT DENSITY FOR $\chi_{BB} = 0.2$

#### PHASE SEPARATION

Direct phase coexistance is used to compute binodals [1]. In order to do so, the simulation box is first centered on the largest cluster. The density is the long axis of the box is then binned and averaged. The resulting distribution is fitted to a hyperbolic profile:

$$\phi(z) = \frac{\rho_L + \rho_G}{2} - \frac{\rho_L - \rho_G}{2} \tanh\left(\frac{z - z_0}{d}\right)$$

Where $z$ is the distance from the center of mass, $\rho_L$ and $\rho_G$ the densities of the two phases, $z_0$ the interface position, and $d$ the interface width. This process yields distinct values for $\rho_L$ and $\rho_G$ even in the single-phase regime. To resolve

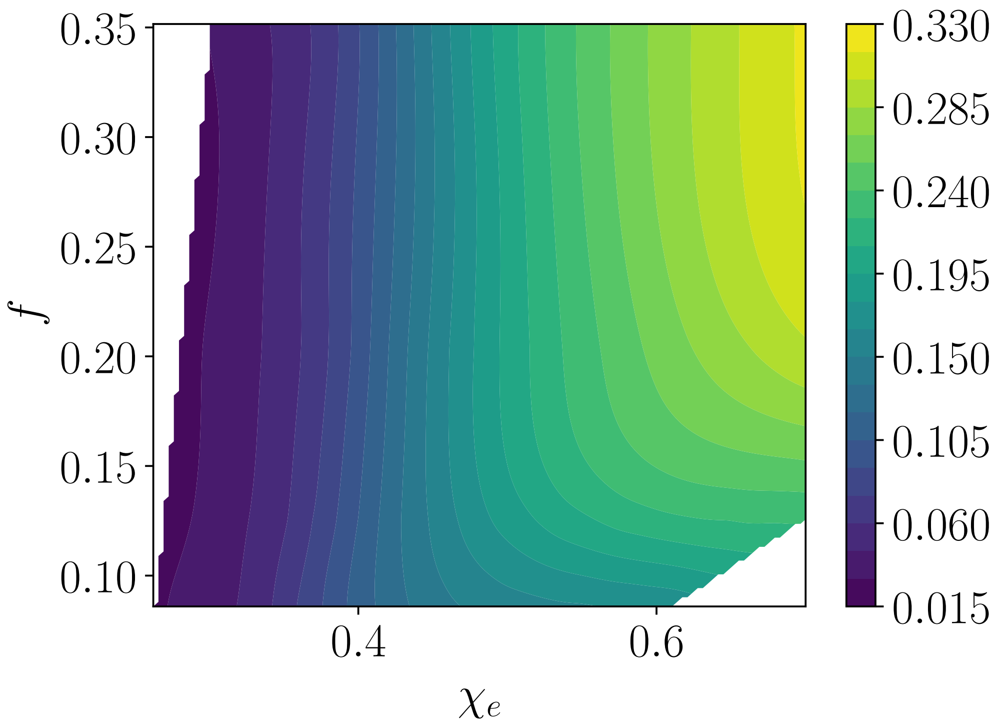


Figure S 1. Mean density of the melt for $\chi_{BB} = 0.2$

*

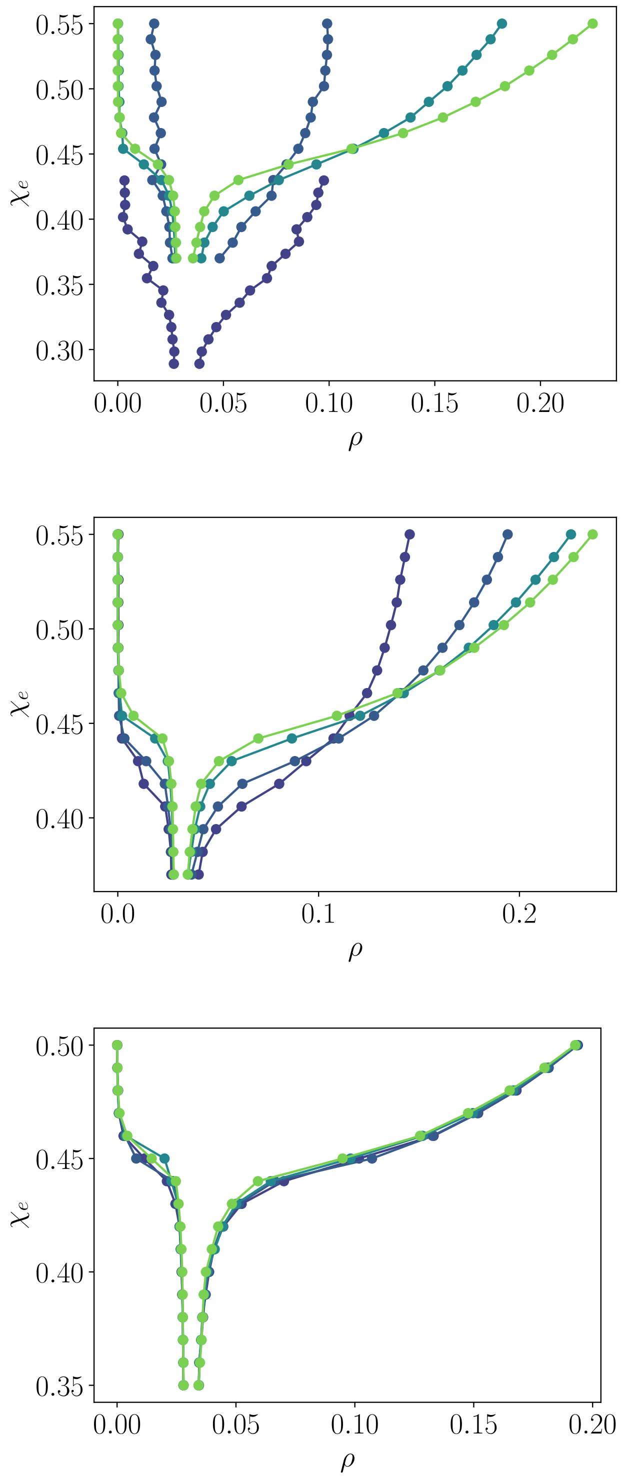


Figure S 2. Binodal curves obtained by fitting hyperbolic profiles to the direct coexistance simulations. Artifacts from the fitting procedure are clearly visibible, as the curve needs to be "convexified". Color indicates values of $f$, from $f = 0.08$ (blue) to $f = 0.35$ (green)

this artifact, a convexification of the binodal curve needs to be considered. For transparency, we report here the fitting results.

[1] G. L. Dignon, W. Zheng, Y. C. Kim, R. B. Best, and J. Mittal, enSequence determinants of protein phase behavior from a coarse-grained model, PLOS Computational Biology **14**, e1005941 (2018), publisher: Public Library of Science.
